# ZZ Top: faster and more adaptive Z chromosome evolution in two Lepidoptera

**DOI:** 10.1101/2020.06.13.142711

**Authors:** Andrew J. Mongue, Megan E. Hansen, James R. Walters

## Abstract

The rate of divergence for Z or X chromosomes is theoretically predicted to be greater than autosomes, but the possible explanations for this pattern vary, as do empirical results from diverse taxa. Even among moths and butterflies (Lepidoptera), which generally share a single-origin Z chromosome, the handful of available studies give mixed support for faster or more adaptive evolution of the Z chromosome, depending on the species assayed. Here, we examine the molecular evolution of Z chromosomes in two additional lepidopteran species: the Carolina sphinx moth and the monarch butterfly, the latter of which possesses a recent chromosomal fusion yielding a segment of newly Z-linked DNA. We find evidence for both faster and more adaptive Z chromosome evolution in both species, though this effect is strongest in the neo-Z portion of the monarch sex chromosome. The neo-Z is less male-biased than expected of a Z chromosome, and unbiased and female-biased genes drive the signal for adaptive evolution here. Together these results suggest that male-biased gene accumulation and haploid selection have opposing effects on long-term rates of adaptation and may help explain the discrepancies in previous findings as well as the repeated evolution of neo-sex chromosomes in Lepidoptera.

## Introduction

Explaining the patterns of genetic variation in natural populations is a foundational goal of population genetics. In the most basic terms, variation is shaped by either selective or random (*i*.*e*. neutral) processes. But beneath this simplicity, dynamics quickly become more complicated. For example, the efficiency of selection relative to drift depends on the effective population size of the genes in question (Ohta 1992). Simple census population size is often a poor proxy for the effective population size, as historical population size changes have long-lasting effects (Tajima 1989). Also, different parts of the genome may have different population sizes due to either differences in ploidy or conditional limitations on expression. For organisms with chromosomal sex determination, the sex chromosomes present a particularly complex confluence of the above processes (Wilson Sayres 2018).

Relative to the rest of the genome, sex chromosomes have smaller population sizes, occurring at either half (in the case of the Y or W) or two thirds (in the case of the X or Z) the frequency of autosomes. Evolution of the Y and W is thought to be driven mainly by the lack of recombination on these chromosomes, leading to the degeneration of all but the most essential genes in many cases (Charlesworth and Charlesworth 2000; Bachtrog 2013). X and Z chromosomes, however, maintain a large set of functional genes in spite of their lower population size. This reduced population size should decrease the efficiency of selection and increase genetic drift (Vicoso and Charlesworth 2009). However, because the X/Z is haploid in one sex, new mutations may be more exposed to selection than on autosomes, causing increased rates of adaptation (Rice 1984; Charlesworth et al. 1987). Both increased drift and increased selection may lead to more rapid rates of molecular evolution on the X/Z relative to autosomes, a phenomenon called “Faster-X” (or, likewise, “Faster-Z”). Although increased divergence of sex chromosomes has been observed repeatedly in diverse taxa, discerning between drift or selection as the primary cause of this Faster-X pattern remains an outstanding challenge, and potentially depends on which sex is heterogametic (Baines et al. 2008; Meisel and Connallon 2013; Kousathanas et al. 2014; Hayes et al. 2020).

A further complication for assessing Faster-X is sex-biased selection on the sex chromosomes. Because of hemizygosity in males, the X spends more time in females than males (and vice versa for the Z). This inequality is predicted to cause an accumulation of sex-biased genes on the sex chromosomes (Rice 1984; Chapman et al. 2003), a pattern that has been observed repeatedly for both male and female heterogametic species (e.g. flies and mice: Meisel et al. 2012; birds: Wright et al. 2012; and butterflies and moths: Mongue and Walters 2017). Genes with sex-biased expression carry their own evolutionary complexities. Because selection can only act on expressed phenotypes, sex-limited genes should be shielded from selection half the time, and thus experience increased divergence due to drift (Dapper and Wade 2016). Sex-specific selection may also influence evolutionary rates and empirical studies indicate sex-biased genes, particularly male-biased genes, evolve faster than unbiased genes, likely due to sexual selection (Grath and Parsch 2016). Thus, an overabundance of sex-linked male-biased genes could in itself cause faster-X, but this scenario is arguably more likely for faster-Z, because Z chromosomes tend to be masculinized while X chromosomes are feminized (Walters and Hardcastle 2011; Meisel et al. 2012; Wright et al. 2012; Mank et al. 2014). However, if the faster-X effect primarily reflects increased adaptation due to ploidy of expression, then it will be particularly pronounced for genes expressed primarily in the heterogametic sex (Baines et al. 2008). Thus faster-Z adaptation should be most apparent for female-biased genes (Parsch and Ellegren 2013; Sackton et al. 2014).

In the better-studied X chromosome systems, a faster-X effect is commonly found, and evidence for more adaptive evolution tends to be associated with species with larger effective population sizes (typically invertebrates, reviewed in Meisel and Connallon 2013). Z chromosome systems are less well-studied, with available results coming mostly from birds. These studies indicate avian Z-linked genes diverge faster primarily due to increased genetic drift, not adaptation (Mank et al. 2009; Wang et al. 2014; Wright et al. 2015; Xu et al. 2019; Hayes et al. 2020), though one study did show increased adaptive divergence of gene expression on the Z (Dean et al. 2015). If larger effective population sizes yield greater faster-X adaptation, then the strongest test for adaptive Z evolution may come from ZW systems with large natural populations, like insects.

Butterflies and moths (Lepidoptera) are the one of the oldest and most diverse female-heterogametic groups and lepidopteran species are routinely observed to have effective population sizes orders of magnitude larger than most vertebrates (Mongue et al. 2019). In spite of these large population sizes, evidence for a lepidopteran faster-Z effect is mixed, with one study finding faster rates of evolution on the Z (Sackton et al. 2014) and two others not (Rousselle et al. 2016; Pinharanda et al. 2019). Likewise, evidence for more adaptation is conflicting, with two of the previous studies finding more adaptation on the Z (Sackton et al. 2014; Pinharanda et al. 2019) and the third finding the opposite: an increase in purifying selection (Rousselle et al. 2016). These contradictory results are particularly baffling given that all Lepidoptera are thought to share a single-origin Z chromosome (Fraïsse et al. 2017) and high levels of synteny (*i*.*e*. conserved gene order) across their phylogeny (Ahola et al. 2014; Davey et al. 2016; Kanost et al. 2016). In other words, the Z chromosome is substantially conserved across taxa; thus, differences in observed molecular evolution may be attributable to a mixture of methodology and lineage-specific effects (*e*.*g*. mating systems skewing effective population sizes). Here, we combine genome-wide polymorphism and divergence data with gene expression analysis in a pair of distantly related Lepidoptera to better understand whether and why the Z chromosome evolves faster than autosomes. We take advantage of robust sequencing data in the Carolina sphinx moth, *Manduca* sexta, and the recent discovery of a neo-Z chromosome in the monarch butterfly, *Danaus plexippus* (Mongue et al. 2017), to examine how newly sex-linked sequence evolves.

## Methods

### Data sources

We intersected population genomic data with sex-specific expression data using published datasets for two Lepidoptera: the Carolina sphinx moth, *Manduca sexta*, and the monarch butterfly, *Danaus plexippus*.

For *M. sexta*, counts of polymorphisms came from whole-genome resequencing of 12 North Carolinian males and divergence came from one sequenced *Manduca quinquemaculata* male (Mongue et al. 2019) aligned to the *M. sexta* reference genome (Kanost et al. 2016). Gene expression levels were obtained from a large RNA-seq dataset that contains numerous stage- and tissue-specific samples (Cao and Jiang 2017). However, sex was not recorded for many of these samples, so we limited our analysis to tissues where comparable male and female data were available: adult heads and antennae, as well as adult and pupal testes and ovaries.

For the monarch butterfly, polymorphisms came from a large-scale resequencing project (Zhan et al. 2014), from which we selected 12 males from the North American migratory population; divergence data came from sequencing of a *Danaus gilippus* male from the same dataset. RNA-seq data came from Illumina sequencing of transcripts from male and female adult butterfly heads, midguts, thoraces, and gonads (ovaries and testes), each sampled in triplicate. Read counts were initially quantified with RSEM (Li and Dewey 2011), then normalized to FPKMs with Trinity using a TMM scaling factor (Grabherr et al. 2011). The three replicates were averaged to give a single expression value per tissue and sex.

For each gene in the genome, we used the SNP-calling pipeline described in Mongue *et al*. (2019). Briefly, we took Illumina resequencing data through the *Genome Analysis Toolkit* best practices pipeline for SNP-calling (McKenna et al. 2010) to generate a set of high-quality variants. We classified each single nucleotide variant as synonymous or non-synonymous using custom databases in *SNPeff* (Cingolani et al. 2012) and normalized variant counts by the number of non-synonymous or synonymous sites in each gene, using custom R scripts to annotate and sum the degeneracy of each amino acid coding site (R Core Team 2017).

### Assignment of sex linkage

Z-linkage in *D. plexippus*, including the presence of a neo-Z segment, was previously characterized using a combination of synteny with other Lepidoptera and differential coverage between male and female sequencing data (Mongue et al. 2017). Z-linkage in *M. sexta* has also been previously assessed, though only via synteny with *Bombyx mori* (Kanost et al. 2016). To directly assess Z-linkage via sex-specific sequencing depth, we generated ∼16x coverage Illumina sequencing from a female *M. sexta* and compared coverage with one of the male samples (S35) having a comparable sequencing depth. For each sample, we used *BEDtools* to calculate per-base coverage across the genome and also the median coverage for each scaffold (Quinlan and Hall 2010). Scaffold medians were normalized by dividing by the mean of all medians for that sample. Then, we assessed Z linkage of each scaffold by taking the log2 of the ratio of male:female normalized coverage for each scaffold. Under this metric, autosomal scaffolds cluster around 0 while Z linked scaffolds group around 1.

### Assessment of Sex-bias

To evaluate sex-bias in expression, we calculated the specificity metric (SPM) for male versus female expression for each annotated gene in the genome (Kryuchkova-Mostacci & Robinson-Rechavi 2017). We first summed FPKM (fragments per kilobase of transcript per million mapped reads) in each sex and divided by the number of replicates for that tissue in that sex to obtain a mean value for each sex and tissue combination. With these values we calculated SPM as the square of expression in one sex divided by the sum of squared expression in both sexes. This calculation gave us a specificity value ranging from 0 to 1, inclusive, indicating what proportion of a given gene’s expression was unique to one sex. As implemented here, an SPM = 1 indicates completely female-specific expression, SPM = 0 indicates male-specific expression, and SPM = 0.5 reflects equal expression between the sexes.

We sought to make our methodology comparable to existing faster-Z studies, which have used fold-change in expression to delineate sex-biased genes. In those analyses, sex-bias cut-offs are typically 1.5x difference in expression between males and females (e.g. in Pinharanda et al. 2019). This difference corresponds to a 70-30 bias in SPM. Thus, we took female-biased genes to be those with SPM > 0.7 in females, male-biased genes with SPM < 0.3 in females, and unbiased genes to be those that fell within (0.3, 0.7). We further verified that our results were robust to cutoff thresholds by re-analyzing the sex-bias data using stricter 85-15 bias. As these results were qualitatively the same as the 70-30 cutoffs, we only present the former in the main text and the latter in the supplement.

We have shown previously that both the *D. plexippus* and *M. sexta* Z chromosomes are masculinized based on distributions of genes encoding sperm proteins (Mongue and Walters 2017), but this RNA-seq expression dataset affords the opportunity to validate those results with a more complete set of sex-biased genes. As such, after classification of sex-biased genes, we used a set of Χ^2^ tests of independence to assess whether or not the proportion of sex-biased genes differed between the autosomes and (neo-)Z chromosomes.

Finally, as Z chromosomes spend more time in males than females and males are often thought to have a higher variance in reproductive success than females, it is possible that the effective population size of the Z is substantially smaller than its census size (Vicoso and Charlesworth 2009). To investigate this possibility, we compared neutral variation between the Z and autosomes in our two study species. In brief, we used a series of custom R scripts to parse out putatively neutral (four-fold degenerate) sites across the genome, and used the population genomics tool ANGSD (Korneliussen et al. 2014) to estimate heterozygosity (Watterson’s θ) separately for all four-fold degenerate sites on the Z and autosomes. We then took the ratio of the mean per-site heterozygosity of the two regions as the difference in effective population size.

### Statistical analysis of molecular evolution

Divergence and polymorphism rates are compound ratios with asymmetric bounds (i.e. both range from 0 to infinity) and are thus not normally distributed. As such, we analyzed molecular evolution with a series of non-parametric tests. Initially, we tested for a faster-Z effect by comparing the scaled rate of divergence (dN/dS) of autosomal and Z-linked genes using Kruskal-Wallis tests with either 1 degree of freedom in *M. sexta* or 2 degrees of freedom in *D. plexippus* to account for 3 potential classes of linkage (autosomal, ancestral Z, and neo-Z). With this distinction established, we assessed the effect of sex-biased gene expression on rates of evolution with another set of Kruskal-Wallis tests to determine if there was an effect of sex-bias generally. In the case of significant results, pair-wise post-hoc differences were investigated with a Nemenyi test. Equivalent tests were performed for polymorphism data as well. Finally, we synthesized the polymorphism and divergence data to calculate α, the proportion of substitutions driven by adaptive evolution. Specifically, we used a modified calculation of the neutrality index (NI) to correct for the bias inherent in a ratio of ratios (Stoletzki and Eyre-Walker 2011) for each class of genes to give us a point-estimate of α (= 1 – NI) summed across genes within a bias class and linkage group. We assessed significance via a permutation test framework, as in Mongue *et al*. (2019). That is, we compared evolution of two gene classes, calculated the point-estimate α for each, then took the absolute value of the difference of these estimates as our permutation test statistic. Next, we combined the two gene sets and randomly drew two permuted classes of sizes equal to the true classes without replacement. We calculated the absolute difference in α for these two random gene sets and repeated this for 10,000 permutations. In doing so, we built a distribution of differences in point estimates of α that could be expected by chance alone. We then compared our true value to this distribution and took the p-value to be the proportion of times we observed a greater value in the permuted distribution than the true value. All of these analyses were completed with custom R scripts in R version 3.4 (R Core Team 2017).

## Results

### Assignment of sex-linkage in *Manduca sexta*

Based on synteny in previous analyses, 27 scaffolds were already annotated as Z-linked in the *M. sexta* assembly (Kanost et al. 2016). By using new resequencing data from a male and female, we validated and updated our knowledge of sex-linkage in this moth. We restricted our coverage analysis to scaffold above the genome N90 (45Kb) to avoid coverage differences that could arise by chance on short sequences. Using this metric, we have recovered all previously annotated 27 scaffolds as male-biased in coverage and thus Z-linked. Additionally, we identified another 9 scaffolds as Z-linked, totaling 2.1Mb and containing 43 annotated genes. Seven of these scaffolds were previously unassigned (due to unclear orthology); the other two had previously been assigned to autosomes but are clearly Z-linked by coverage. The updated linkage information is included as a supplementary datatable.

### Sex-bias on the Z chromosomes

Based on the assignment of sex-biased genes from the RNA-sequencing data, the gene-content differs between the Z and autosomes in both the Carolina sphinx moth (Χ^2^_2_ = 47.37, p = 5.2*10^−11^) and monarch butterfly (Χ^2^_2_ = 30.04, p = 3.0*10^−7^). In both species, this difference comes from an excess of male-biased genes on the Z chromosome, as well as a paucity of female-biased genes on the *Manduca* Z and unbiased genes on the *Danaus* ancestral Z (Table 1). These results hold for both these cutoffs for sex bias and for stricter criteria (85-15 SPM, see supplement). It is worth noting that the male bias in expression on the Z chromosome is not the result of dosage effects, as both *M. sexta* and *D. plexippus* have been shown generally to have sex-balanced expression on the Z chromosome (Smith et al. 2014; Gu and Walters 2017).

**Table 1.** Sex bias of the Z chromosomes in the two species studied with raw counts and proportions in parentheses. In both species, composition of the Z differs from composition of the autosomes due to an increased proportion of male-biased Z-linked genes (based on Χ^2^ p-values < 1.0*10^−6^; note that this significant result holds in monarchs whether the Z is considered as one category or two (i.e. neo and ancestral)). The sphinx moth Z is depleted for female-biased genes, while the monarch (ancestral-)Z is depleted for unbiased genes.

|  | Carolina sphinx moth |  | Monarch butterfly |  |  |
| --- | --- | --- | --- | --- | --- |
|  | Autosomes | Z | Autosomes | Ancestral Z | Neo-Z |
| Male-biased | 2477 (0.21) | 177 (0.34) | 4721 (0.35) | 279 (0.47) | 184 (0.39) |
| Unbiased | 7219 (0.63) | 295 (0.56) | 7529 (0.56) | 278 (0.46) | 243 (0.52) |
| Female-biased | 1856 (0.16) | 55 (0.10) | 1248 (0.09) | 44 (0.07) | 41 (0.09) |

### Rates of divergence

We found that the Z chromosome has higher scaled divergence than the autosomes in both species, the Carolina sphinx moth (Χ^2^_1_ = 6.89, p = 0.009, Figure 1A) and the monarch butterfly (Χ^2^_2_ = 9.72, p = 0.008). For monarchs, we further classified the Z into the ancestral- (*i*.*e*. long-term sex-linked) and neo-Z (the Z sequence resulting from a milkweed-butterfly-specific autosomal fusion). Based on the significant chromosomal linkage effect, we conducted post-hoc testing and found that the signal for faster-Z evolution comes primarily from the neo-Z, which diverges distinctly faster than the autosomes (p = 0.006, Figure 1D) and marginally faster than the ancestral Z (p = 0.048). The ancestral Z was not faster evolving than the autosomes (p = 0.99).

**Figure 1.**
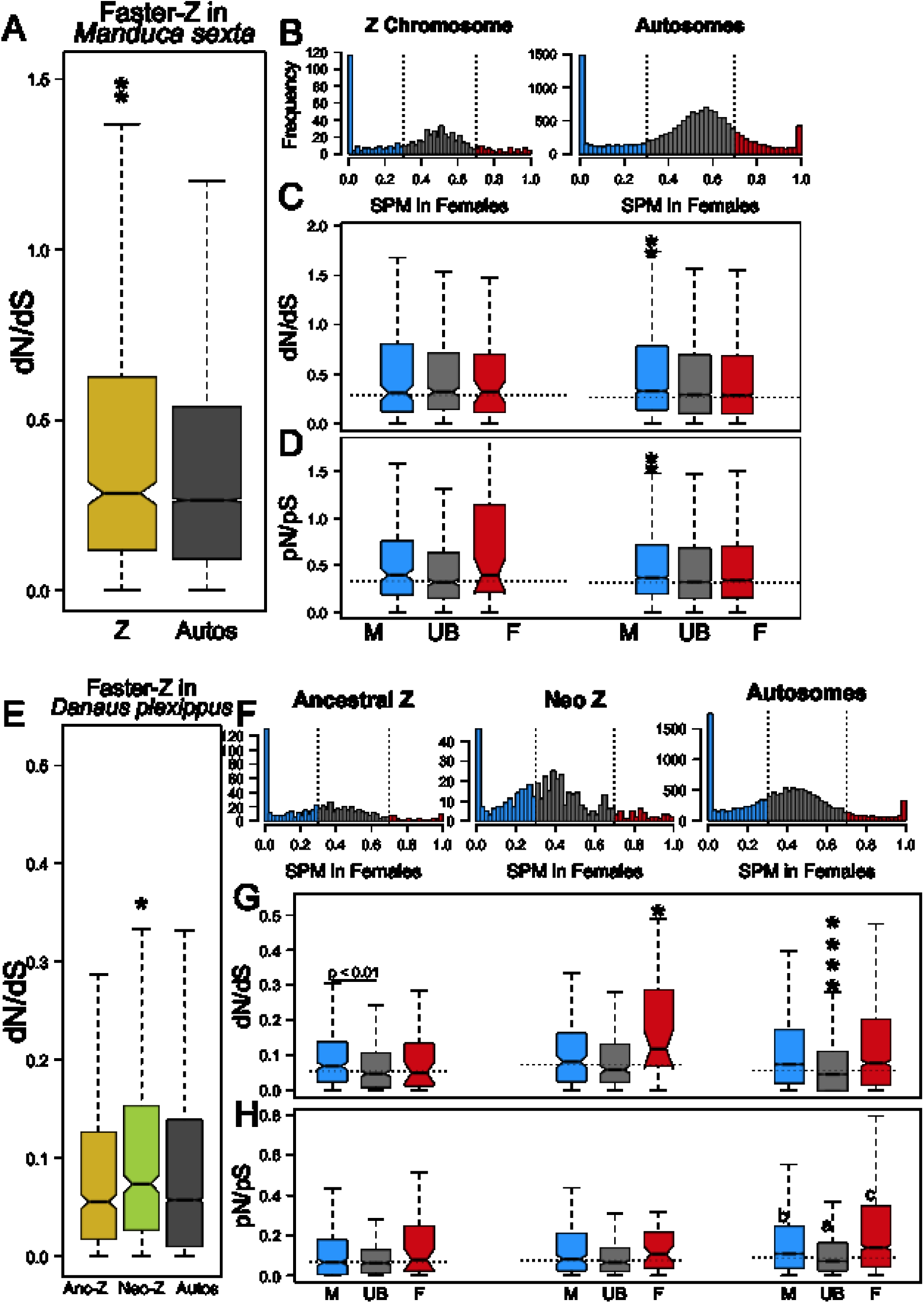
Faster-Z evolution in Carolina sphinx moths and monarchs. Throughout, asterisks represent statistical differences of one group from all others to which it is compared, with the number of asterisks indicating the level of significance (* < 0.05, ** <0.01, *** <0.001, etc.). Horizontal lines with significance annotations are given for significant pairwise differences. **A**. The Z evolves faster than the autosomes in *Manduca sexta*. **B**. The distributions of sex-bias for both z-linked (left) and autosomal (right) genes are plotted with dashed lines to indicate the traditional cutoff points for sex-bias analysis. Bias is plotted such that higher SPM values are more female biased in expression, while values closer to 0 are male-biased. **C**. Rates of divergence for genes in each sex-bias class (M: male-biased, UB: unbiased, F: female-biased). In sphinx moths, only autosomal genes show differences between rates of evolution of genes with different sex-bias. **D**. Likewise, male-biased genes have higher pN/pS than on other bias classes, but only on the autosomes. **E**. The neo-Z is the source of a faster-Z signal in monarch butterflies. **F**. Again we plot distributions of sex-bias categories for genes on the ancestral Z (left), neo-Z (middle), and autosomes (right). **G**. In monarchs, male-biased genes evolve more quickly on the ancestral Z. Female biased genes evolve more quickly on the neo-Z, and unbiased genes evolve more slowly on the autosomes. **H**. Finally, sex-biased genes hold different levels of polymorphism on the autosomes, with unbiased genes having the lowest pN/pS, followed by male-biased, then female-biased with the highest (graphically represented as a < b < c).

In Carolina sphinx moths, sex-biased expression did not impact divergence rates on the Z chromosome (Χ^2^_2_ = 1.12, p = 0.571, Figure 1C). On the autosomes however, there was a clear effect of sex-biased expression (Χ^2^_2_ = 26.26, p = 1.98*10^−6^). Post-hoc testing revealed this to be driven largely by male-biased genes, which had higher divergence rates than unbiased (p = 8.1*10^−6^) or female-biased genes (p = 4.2*10^−5^). Female-biased genes do not evolve at a different rate than unbiased genes (p = 0.63).

In monarchs, like sphinx moth, sex-biased expression affected evolutionary rates of autosomal loci (Χ^2^_2_ = 249, p < 1.0 * 10^−10^, Figure 1F). Unlike sphinx moths however, the effect of sex-bias did not differ between sexes. Both male-biased (p < 1.0*10^−10^) and female-biased genes (p < 1.0 * 10^−10^) evolve faster than unbiased genes according to post-hoc testing, though male-biased and female-biased genes did not evolve differently from each other (p = 0.75).

Considering the monarch Z chromosome, both the ancestral (Χ^2^_2_ = 9.99, p = 0.007) and neo (Χ^2^_2_ = 11.85, p = 0.003) segments showed a sex-bias effect. For the ancestral Z, this difference is driven solely by faster evolution of male-biased genes compared to unbiased genes (p = 0.005); evolutionary rates of female biased genes did not differ significantly from the unbiased nor male-biased genes on the ancestral Z. On the neo-Z, female-biased genes evolve faster than both male-biased (p = 0.044) and unbiased genes (p = 0.002); divergence of male-biased genes did not differ from unbiased on the neo-Z.

### Rates of polymorphism

Unlike with divergence, we found that that sex-linkage did not strongly impact rates of polymorphism in *M. sexta* (Χ^2^_1_ = 2.57, p = 0.110) However, sex-biased expression did impact rates of polymorphism (Χ^2^_2_ = 43.45, p = 3.7 * 10^−10^). Here again, male-biased genes showed increased variation compared to unbiased genes (p = 1.4 * 10^−10^) and female-biased genes (p = 0.002). Female-biased and unbiased genes did not significantly differ from each other (p = 0.14).

In monarchs, sex linkage strongly impacted polymorphism (Χ^2^_2_ = 34.18, p = 38 * 10^−8^). Both the ancestral Z (p = 3.9 * 10^−7^) and neo-Z (p = 0.02) had lower rates of polymorphism than the autosomes, but the two portions of the Z did not differ from each other (p = 0.27). Sex-biased expression did not significantly affect polymorphism on either part of the Z (ancestral: Χ^2^_2_ = 2.70, p = 0.259; neo: Χ^2^_2_ = 5.75, p = 0.06). In contrast, autosomal genes did show an effect of sex-bias, with female-biased genes showing the highest rates of polymorphism, higher than male-biased (p = 1.8*10^−10^) or unbiased genes (p < 1.0 * 10^−10^); male-biased genes had elevated rates of polymorphism compared to unbiased genes (p < 1.0 * 10^−10^).

### Rates of adaptive evolution

The variable patterns of divergence and polymorphism observed for sex-linked and sex-biased loci may reflect differing rates of adaptive evolution among these groups of genes. To examine this, we estimated the proportion of adaptive substitutions (α) for each gene-class, first contrasting the Z versus autosomes as a whole, and subsequently further partitioning loci by sex-biased expression.

In *M. sexta*, the Z overall showed more adaptive evolution than the autosomes (p = 0.039). Adaptation of male-biased (p = 0.340) and female-biased genes (p = 0.812) did not differ based on genomic location, but genes with unbiased expression showed higher rates of adaptive evolution (α) on the Z chromosome than the autosomes (p = 0.007; Figure 2A).

**Figure 2.**
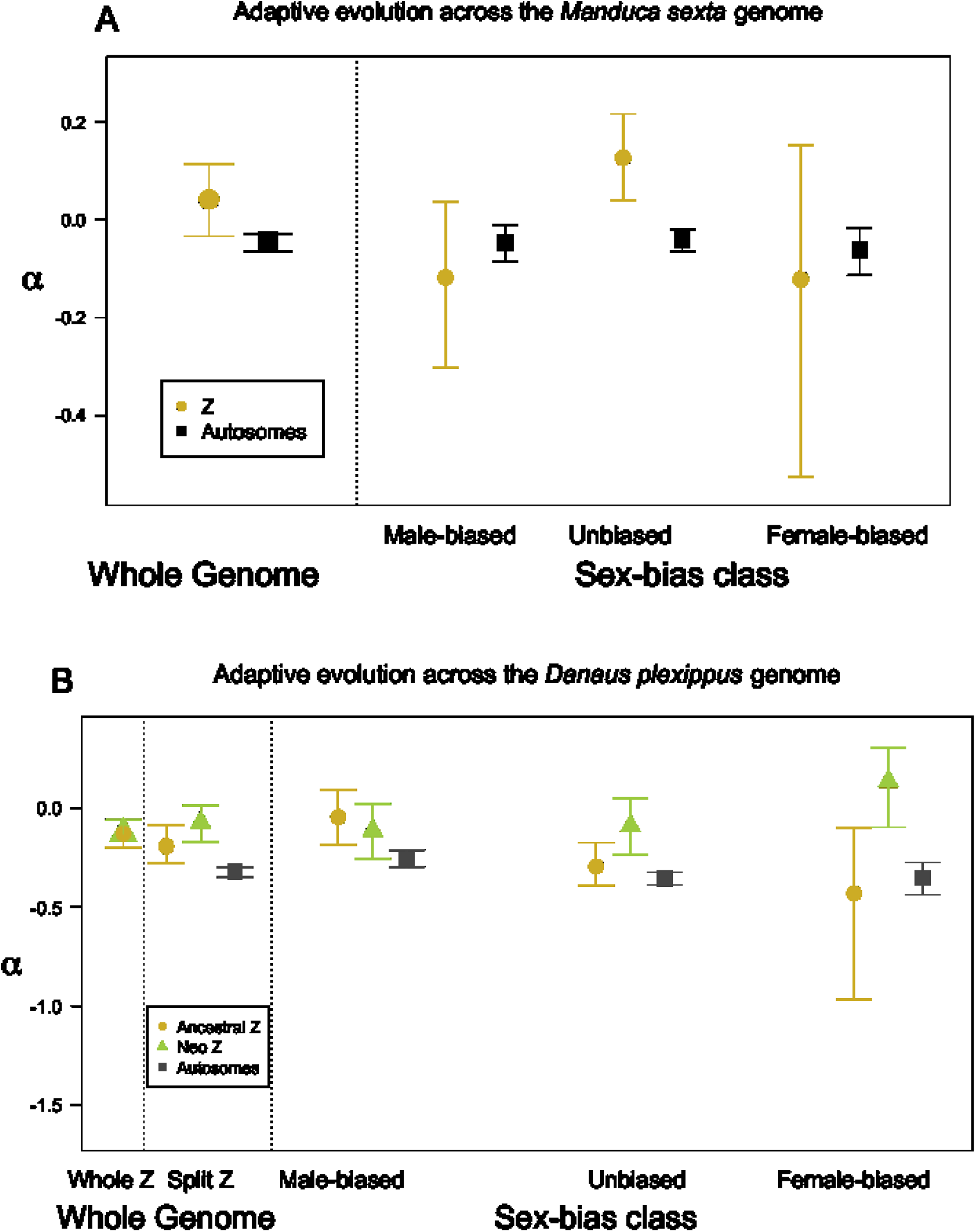
Adaptive evolution across the genomes of the two Lepidoptera considered in this study. In each panel coarse-scale comparison of the Z chromosome to autosomes are plotted left of the dotted lines. Points are the point estimate of the α statistic and error bars represent 95% confidence intervals for each point estimate obtained by parametric bootstrapping. In the Carolina sphinx moth (**A**), the Z evolves more adaptively than the autosomes overall (left of dash). This pattern appears to be driven by unbiased genes (right of dash). In the monarch butterfly (**B**), the whole Z is more adaptively evolving than the autosomes (leftmost), and both the ancestral and neo-segments show elevated α compared to the autosomes (middle). For the ancestral Z, male-biased genes drive the increase in adaptation; in contrast, unbiased and female-biased genes are more adaptively evolving on the neo-Z.

In monarchs, the Z also exhibited increased rates of adaptation compared to autosomes (p = 0.0004; Figure 2B, left). Considered separately, both the ancestral and neo-Z segments evolved more adaptively than the autosomes (ancestral-Z vs. autosomes: p = 0.0338, neo-Z vs. autosomes: p = 0.0005). The neo-Z segment trended towards more adaptive evolution than the ancestral Z, but not strongly (p = 0.079). Turning to sex-bias, we found that male-biased genes did not evolve differently across the genome (autosomal vs. neo-Z p = 0.318, autosomal vs. ancestral Z p = 0.092, ancestral vs neo-Z p = 0.500). In contrast, female-biased genes evolved more adaptively on the neo-Z than the autosomes (p = 0.0474) or ancestral Z (p = 0.008). Additionally, ancestrally Z-linked female-biased genes did not evolve differently than their autosomal counterparts (p = 0.539). Furthermore, unbiased genes on the neo-Z showed greater rates of adaptation than unbiased genes on the autosomes (p = 0.018) or ancestral Z (p = 0.048).

### The effective population size of the Z chromosome

Under the simplest biological conditions, we expect the ratio of population sizes of Z:Autosomes to be 0.75 (because males and females each make up half of the population (Wilson Sayres 2018)). We found for *Manduca sexta* that in practice this ratio is much smaller Ne_Z_:Ne_A_ = 0.44. For *Danaus plexippus*, the difference in population sizes is less skewed, Ne_Z_:Ne_A_ = 0.66. Intriguingly, this difference is not uniform across the monarch Z. The ancestral portion of the Z has a lower population size, Ne_Z_Anc_:Ne_A_ = 0.58, but the neo-Z holds essentially as much diversity as the autosomes, Ne_Z_Neo_:Ne_A_ = 0.98.

## Discussion

### New evidence for a faster-Z

While previous evidence for faster-Z evolution in Lepidoptera has been mixed, we found that the Z chromosome is faster evolving (*i*.*e*. has a higher dN/dS ratio) than the autosomes in both *Manduca sexta* and *Danaus plexippus*. At first pass, our results seemingly align with a report of faster-Z evolution in silkmoths (Sackton et al. 2014), but are contrasting with other studies in butterflies (Rousselle et al. 2016; Pinharanda et al. 2019). However, a more nuanced consideration indicates some congruence with both sets of studies. Monarchs show an overall fast Z, but this result is driven by the newly sex-linked neo-Z portion of the monarch evolving faster than the autosomes. Considering only the ancestral portion, which is homologous to the Z of the butterflies previously studied, there is no evidence for a faster ancestral-Z in monarchs. However, evidence for higher rates of adaptive evolution (α) on the Z is less ambiguous in our insects; both *Manduca* and *Danaus* showed overall more adaptation for Z-linked genes, as reported in *Bombyx*.

Beginning with the simpler case of *Manduca*, we found that increased adaptation on the Z chromosome is driven by genes with unbiased (*i*.*e*. equal) expression in the two sexes. These genes will be expressed in the haploid state in females and thus should experience more efficient selection than unbiased genes on the autosome (which are always diploid expressed). Female-biased genes should theoretically follow this pattern as well, but the lack of a clear signal might be partially attributable to the relatively small number of female-biased genes on the Z, which reduces power to detect differences in adaptive evolution. Moreover, the effective population size of the *Manduca* Z compared to the autosomes is much lower than the neutral expectation (0.44, as opposed to 0.75). With such a decrease in the population of Z chromosomes, selection is predicted to be less efficient (Vicoso and Charlesworth 2009) and may further limit the adaptive evolution of female-biased genes.

*Danaus* presents a more complicated case, owing to a Z-autosomal fusion in this genus (Mongue et al. 2017). Intriguingly, it is the neo-Z that best fits with theoretical predictions for adaptive Z evolution. Increased adaption (compared to the autosomes) is concentrated in unbiased and female-biased genes as expected. It also is worth noting that the neo-Z has an inferred effective population size nearly equal to that of the autosomes (Ne_Z_neo_:Ne_Autos_ = 0.98). This is a perplexing result that cannot be attributed to sequence homology with a neo-W, which if exists at all must be highly divergent from the neo-Z, (Mongue et al. 2017; Gu et al. 2019). Parity in effective population size of the sex chromosomes and autosomes has been attributed to biased sex ratios and/or higher variance in the reproductive success of the heterogametic sex in other taxa (Hedrick 2007; Ellegren 2009). A skewed sex ratio seems unlikely, as only a male-biased population could restore parity to the Z:A ratio. No such dynamics have been observed (on the contrary, another *Danaus* species is known to have male-killing genetic elements (Smith et al. 2016)), and more to the point, this dynamic should affect the ancestral and neo-Z equally. A high variance in female reproductive success could also generate roughly equal effective population sizes of the Z and autosomes and should be more apparent in regions with fewer male-biased genes (*i*.*e*. the neo-Z), but there is currently little ecological data with which to assess this possibility. Whatever the cause of this parity in variation, this observation itself points to different evolutionary dynamics between the ancestral and neo-Z, and implies that selection to remove deleterious variation should be more efficient than on the autosomes for all dominance coefficients of deleterious mutations (Vicoso and Charlesworth 2009).

In contrast to the neo-Z, the *Danaus* ancestral Z has a much lower effective population size compared to the autosomes (0.58), yet it evolves more adaptively than the autosomes overall. This result is driven by increased adaptation among Z-linked male-biased genes. As discussed above, this result points to a difference in evolutionary dynamics between the neo- and ancestral segments. Moreover, this suggests that, in terms of genetic diversity, the difference in census size between the autosomes and sex chromosomes is relatively unimportant in this butterfly. One possible explanation is the unusually strong sexual selection in this species. Female monarchs are highly polyandrous in nature (Pliske 1975), which has been implicated in the elevated adaptation of sperm protein coding genes on the autosomes (Mongue et al. 2019). As the ancestral Z is highly masculinized in gene content, it stands to reason that similarly strong selection may apply to these male-biased genes. Circumstantial evidence for this can be seen in the effective population size ratios of the ancestral Z segment compared to the non-masculinized neo-Z segment.

### Reconciling existing investigations of lepidopteran Z chromosome evolution

Our results most strongly agree with existing work from the silkmoth genus *Bombyx* (Sackton et al. 2014), which found both fast and adaptive Z effects. Efforts in other butterflies have found no fast Z effect. In the case of satyr butterflies, this negative result may be attributable to “noisy” sequence data (*de novo* transcriptome assemblies were used) and potential uncertainty in Z-linkage (which was inferred from sequence homology to another butterfly species) (Rousselle et al. 2016). Separately, in the case of *Heliconius* butterflies, it is worth noting that point estimates for α and dN/dS largely fit predictions for a fast and adaptive Z, but results did not differ significantly between the Z and autosomes thanks to high variance in these estimates, especially on the Z chromosomes (Pinharanda et al. 2019). In this case, the use of a relatively small RNA-sequencing dataset created a smaller dataset of sex-biased genes with which to work, and only 200 of about 700 total Z-linked genes were analyzed.

Nonetheless, this collection of lepidopteran faster-Z studies suggests a phylogenetic signal for Z chromosome evolution. *Bombyx* and *Manduca* are species from sister families of moths (Kawahara and Breinholt 2014) and share patterns of faster and more adaptive Z evolution. Satyrs, *Heliconius*, and *Danaus* butterflies all fall within the family Nymphalidae and show mixed to negative evidence for increased divergence and adaptation on the (ancestral) Z. In other words, there is more agreement for Z chromosome evolution for more closely related species (though insects in general and Lepidoptera specifically are an ancient group, so none of these species are “closely-related” by vertebrate-centric expectations). These observations demonstrate that sex-linkage *per se* does not lead to consistent evolutionary outcomes for the genes involved. Instead faster-Z evolution is likely to depend on the demographic history or degree of sex-bias of the Z chromosomes examined. This is illustrated by the relatively young neo-Z in monarchs, which is not masculinized like the ancestral lepidopteran Z sequences and instead appears comparable to autosomes in the proportion of unbiased and female-biased genes (Mongue and Walters 2017). Intriguingly, the monarch neo-Z fits completely within the theoretical prediction for adaptive faster-Z evolution. It appears to be faster evolving due to increased adaptation of unbiased and female-biased genes that are subject to haploid selection (Charlesworth et al. 1987). These observations present the possibility that faster-Z dynamics may be transient rather than perpetual.

Indeed, two prominent hypotheses for sex chromosome evolution combine to suggest this transient faster-Z dynamic. Adaptive evolution of the sex chromosomes is thought to be driven by the hemizygous expression of some genes in one sex (Charlesworth et al. 1987), but depending on the dominance of gene expression, genes benefitting the opposite sex are predicted to accumulate on that sex chromosome (Rice 1984). As such, if the sex chromosomes change composition over evolutionary time, they may bias towards alleles benefitting the homogametic sex (e.g. male-benefitting, male-biased genes on the Z). Genes with haploid expression (e.g. unbiased or female-biased genes in ZZ/ZW systems), will become less abundant and thus less important to the overall evolution of the chromosome. Moreover, if sexual selection produces high variance in male reproductive success, the effective population size of Z chromosomes can be substantially depressed below the census size, further limiting the role of positive selection on the few unbiased or female-biased left on the Z. Particularly old sex chromosomes should be more likely to experience these effects.

To take this thread of logic to its end, this scenario may also explain the relative abundance of neo-Z chromosomes in Lepidoptera (Nguyen et al. 2013; Nguyen and Paladino 2016; Mongue et al. 2017). The strongly conserved synteny across species implies that small-scale gene trafficking events are rare (but evidence is somewhat contradictory here as well, see: Toups et al. 2011; Wang et al. 2012) and fusion-fission events may be the key source of linkage shuffling in Lepidoptera. For a highly masculinized Z chromosome, a sudden influx of unbiased and female-biased genes onto the Z can create strong positive selection and favor these fused chromosomes, even at initially low frequencies, helping them to escape elimination by drift. Under this paradigm, even the seemingly contradictory findings on Z chromosome evolution can be reconciled as being the product of lineage-specific differences in sex-biased gene content and chromosomal history. If this line of reasoning is accurate, it should be borne out in other Lepidoptera with neo-Z chromosomes. More comparisons of independent neo-Z chromosomes will be needed for this validation.

### Data accessability

*Manduca sexta* whole genome resequencing data can be found on NCBI’s Sequence Read Archive with the following accessions: SRP144217, PRJNA639154. *Danaus plexippus* RNA sequencing can be found with PRJNA522622. The *M. sexta* expression data can be found as a supplementary table in Cao and Jiang (2017), https://doi.org/10.1186/s12864-017-4147-y. The *D. plexippus* sequencing data can be found in Zhan et al. (2014), https://doi.org/10.1038/nature13812. Please see the supplement to this manuscript for specific samples used here.

## Acknowledgements

This project was funded by the NSF DDIG (DEB-1701931). The authors wish to acknowledge Wesley Mason and Michael Hulet and the rest of the Information and Telecommunication Technology Center (ITTC) staff at the University of Kansas for their support with our high-performance computing. We are continually grateful to Chip Taylor and Ann Ryan from MonarchWatch for access to butterflies and to Clyde Sorenson for logistical and moral support in obtaining sphinx moths. Thanks, as well, to Alex Mackintosh and other frequenters of the Darwin Dance Hall for conversations during the interpretation of these results.

## Notes

### Competing Interest Statement

The authors have declared no competing interest.

