## Supplementary methods for "ZZ Top: faster and more adaptive Z chromosome evolution in two Lepidoptera"

**Samples used for genomic analysis**

Whole genome resequencing for both polymorphism and divergence data came from previously published sources. In an effort to increase repeatability and transparency, we have summarized these samples below.

| Species | Sample | Sex | Published in | Accession |
|---|---|---|---|---|
| *Manduca sexta* | LK5 | F | This manuscript | PRJNA639154 |
| *Manduca sexta* | S32 | M | Mongue et al., 2019 | SRP144217 |
| *Manduca sexta* | S33 | M | Mongue et al., 2019 | SRP144217 |
| *Manduca sexta* | S34 | M | Mongue et al., 2019 | SRP144217 |
| *Manduca sexta* | S35 | M | Mongue et al., 2019 | SRP144217 |
| *Manduca sexta* | S36 | M | Mongue et al., 2019 | SRP144217 |
| *Manduca sexta* | S37 | M | Mongue et al., 2019 | SRP144217 |
| *Manduca sexta* | S38 | M | Mongue et al., 2019 | SRP144217 |
| *Manduca sexta* | S39 | M | Mongue et al., 2019 | SRP144217 |
| *Manduca sexta* | S40 | M | Mongue et al., 2019 | SRP144217 |
| *Manduca sexta* | S42 | M | Mongue et al., 2019 | SRP144217 |
| *Manduca sexta* | S44 | M | Mongue et al., 2019 | SRP144217 |
| *Manduca sexta* | S45 | M | Mongue et al., 2019 | SRP144217 |
| *Manduca quinquemaculata* | Q6 | M | Mongue et al., 2019 | SRP144217 |
| *Danaus plexippus* | Plex_TX_T14 | M | Zhan et al., 2014 | SRP045468 |
| *Danaus plexippus* | Plex_TX_T9 | M | Zhan et al., 2014 | SRP045468 |
| *Danaus plexippus* | Plex_MEX_986 | M | Zhan et al., 2014 | SRP045468 |
| *Danaus plexippus* | Plex_MEX_919 | M | Zhan et al., 2014 | SRP045468 |
| *Danaus plexippus* | Plex_MEX_536 | M | Zhan et al., 2014 | SRP045468 |
| *Danaus plexippus* | Plex_MA_HI032 | M | Zhan et al., 2014 | SRP045468 |
| *Danaus plexippus* | Plex_StMFL_146 | M | Zhan et al., 2014 | SRP045468 |

| | | | | |
|---|---|---|---|---|
| *Danaus plexippus* | Plex_CA_SB146 | M | Zhan et al., 2014 | SRP045468 |
| *Danaus plexippus* | Plex_FLn_StM105 | M | Zhan et al., 2014 | SRP045468 |
| *Danaus plexippus* | Plex_StMFL_109 | M | Zhan et al., 2014 | SRP045468 |
| *Danaus plexippus* | Plex_NJ_116 | M | Zhan et al., 2014 | SRP045468 |
| *Danaus plexippus* | Plex_BOS_HI023 | M | Zhan et al., 2014 | SRP045468 |
| *Danaus gilippus* | Gili_CRC_30 | M | Zhan et al., 2014 | SRP045468 |

**Stricter sex-bias analyses**

We repeated analyses of sex-biased expression with stricter cutoffs (i.e. defining a sex-biased gene as one with 85% of expression in one sex instead of 70% as in the main text). This new threshold decreased the number of genes considered as male- or female-biased and thus necessarily decreased statistical power, but our goal here was to demonstrate that our findings are not contingent on how we define sex-biased genes. The statistical inference is the same as in the main methods, so will not be repeated here for brevity.

**Sex-bias on the Z chromosomes**

The gene-content differs between the Z and autosomes in both the Carolina sphinx moth ($X^2_2 = 36.76$, $p = 1.0*10^{-8}$) and monarch butterfly ($X^2_2 = 17.12$, $p = 2.0*10^{-4}$). In both species, this difference comes from an excess of male-biased genes on the Z chromosome, as well as a small of female-biased genes on the *Manduca* Z and unbiased genes on the *Danaus* ancestral Z (Table S1). These results are in complete agreement with the main text's analysis of sex bias.

| | Carolina sphinx moth | | Monarch butterfly | | |
|---|---|---|---|---|---|
| | Autosomes | Z | Autosomes | Ancestral Z | Neo-Z |
| Male-biased | 1738 (0.15) | 131 (0.25) | 2794 (0.21) | 189 (0.31) | 84 (0.18) |
| Unbiased | 9091 (0.79) | 371 (0.70) | 10049 (0.74) | 391 (0.65) | 369 (0.79) |
| Female-biased | 713 (0.06) | 25 (0.05) | 655 (0.05) | 21 (0.04) | 15 (0.03) |

**Table S1.** Sex bias of the Z chromosomes in the two species studied with a stricter definition of sex-biased expression. In both species, composition of the Z differs from composition of the autosomes due to an increased proportion of male-biased Z-linked genes (based on $X^2$ p-values < $1.0*10^{-3}$; again, this significant result holds in monarchs whether the Z is considered as one category or two (i.e. neo and ancestral)).

**Rates of divergence**

In Carolina sphinx moths, sex-biased expression did not impact divergence rates on the Z chromosome ($X^2_2$ = 0.98, p = 0.614).On the autosomes however, there was a clear effect of sex-biased expression ($X^2_2$ = 27.37, p = $1.14*10^{-6}$). Post-hoc testing revealed this to be driven largely by male-biased genes, which had higher divergence rates than unbiased (p = $2.0*10^{-5}$) or female-biased genes (p = $1.7*10^{-5}$). Female-biased genes trended towards slower evolution than unbiased genes, but not significantly so (p = 0.078).

In monarchs, like sphinx moth, sex-biased expression affected evolutionary rates of autosomal loci ($X^2_2$ = 341, p < 1.0 * $10^{-10}$). Unlike sphinx moths however, the effect of sex-bias did not differ between sexes. Both male-biased (p < $1.0*10^{-10}$) and female-biased genes (p < 1.0 * $10^{-10}$) evolve faster than unbiased genes according to post-hoc testing, though male-biased and female-biased genes did not evolve differently from each other (p = 0.28).

Considering the monarch Z chromosome, both the ancestral ($X^2_2$ = 24.7, p = $4.31*10^{-6}$) and neo ($X^2_2$ = 21.1, p = $2.57*10^{-5}$) segments showed a sex-bias effect. For the ancestral Z, this difference is driven solely by faster evolution of male-biased genes compared to unbiased genes (p = $3.30*10^{-6}$); evolutionary rates of female biased genes did not differ significantly from the unbiased nor male-biased genes on the ancestral Z. On the neo-Z, the relatively small number of female-biased genes (15) lowers our power to detect differences, and the evolution of the remaining female-biased genes cannot be distinguished from that of male-biased (p = 0.96) or unbiased (p = 0.08) genes. Nevertheless, the median dN/dS for strongly female-biased genes on the neo-Z is 0.118, compared to 0.060 for unbiased genes, suggesting that the trend for faster evolution of female-biased genes holds here as well. The key difference with the more strict analyses is that the most strongly male-biased genes evolve faster than unbiased genes on the neo-Z (p = $7.20*10^{-5}$, median dN/dS = 0.121). There are relatively few (84) of these male-biased genes, and, as we argue in the main text, they apparently contribute little to the overall chromosome's evolution, but this finding does suggest that new sex-linkage may be driving rapid evolution of the few male-biased genes on the neo-Z segment.

**Rates of polymorphism**

Although sex-linkage did not affect rates of polymorphism in *Manduca,* sex-biased expression did ($X^2_2$ = 41.85, p = 8.2 * $10^{-10}$). Here again, male-biased genes showed the strongest difference from unbiased genes (p = 4.5 * $10^{-10}$) but were also weakly different from female-biased genes (p = 0.054). Female-biased and unbiased genes did not significantly differ from each other (p = 0.26).

Sex-biased expression did not strongly affect polymorphism on the ancestral Z ( $X^2_2$ = 4.88, p = 0.09); however, under stricter filtering, sex-biased genes showed different patterns of variation on the neo-Z ($X^2_2$ = 7.30, p = 0.03).

In contrast, autosomal genes did show an effect of sex-bias, with female-biased genes showing the highest rates of polymorphism, higher than male-biased ($p = 1.8*10^{-10}$) or unbiased genes ($p < 1.0 * 10^{-10}$); male-biased genes had elevated rates of polymorphism compared to unbiased genes ($p < 1.0 * 10^{-10}$). This difference is driven by a decrease of polymorphism of unbiased genes compared to male-biased genes (p = 0.02), but not female-biased genes (p = 0.25). As mentioned above, it is worth remembering that the paucity of strongly female-biased genes lowers our power to detect differences, but the median rates of polymorphism (pN/pS) are lower for both female-biased (0.051) and unbiased genes (0.069) compared to male-biased genes (0.121) on the neo-Z. This pattern is consistent with more efficient purifying selection on genes with at least some hemizygous expression.

**Rates of adaptive evolution**

Our calculation of α quickly loses power with smaller sample sizes; nevertheless, we recalculated differences in adaptation to confirm that patterns did not drastically change with the stricter thresholds for sex-bias. Starting with *Manduca*, adaptation of male-biased (p = 0.198) and female-biased genes (p = 0.796) did not differ based on genomic location, but genes with unbiased expression still showed higher rates of adaptive evolution on the Z chromosome than the autosomes (p = 0.011). These results are in complete agreement with the less strict thresholds.

Turning to *Danaus*, we found that male-biased genes did not evolve differently across the genome (autosomal vs. neo-Z p = 0.891, autosomal vs. ancestral Z p = 0.083, ancestral vs neo-Z p = 0.327). With fewer female-biased genes, we no longer recovered more adaptive evolution on the neo-Z than the autosomes (p = 0.141) or ancestral Z (p = 0.0023). Additionally, ancestrally Z-linked female-biased genes did not evolve differently than their autosomal counterparts (p = 0.086). Furthermore, unbiased genes on the neo-Z still showed greater rates of adaptation than unbiased genes on the autosomes (p = 0.0002) or the ancestral Z (p = 0.021).

In conclusion, considering only strongly sex-biased genes does not change our conclusions about divergence, polymorphism, or adaptive evolution.

**A note on the updated *Manduca* linkage file**

Included with this manuscript is a dataframe offering updated linkage information, polymorphism, divergence (with *M. quinquemaculata*), and sex-specificity data. For sex linkage, another categorical variable (“ZorA”) is appended for ease of analysis. However, for sex-biased expression, no firm category is included, as, per the point of this supplement, the researcher may wish to investigate how molecular variation changes (or does not) with differing cutoff specificities of SPM.
